# Orca vowels and consonants: convergent spectral structures across cetacean and human speech

**DOI:** 10.64898/2026.02.27.708287

**Authors:** Gašper Beguš, Marla Holt, Brianna Wright, David F. Gruber

**Affiliations:** Department of Linguistics, University of California, Berkeley, United States; Project CETI, New York, United States and Roseau, Dominica; Conservation Biology Division, Northwest Fisheries Science Center, National Marine Fisheries Service, National Oceanic and Atmospheric Administration, Seattle, Washington, USA; Pacific Biological Station, Fisheries and Oceans Canada, Nanaimo, British Columbia, Canada; Department of Natural Sciences, Baruch College and The Graduate Center, City University of New York, NY United States

## Abstract

The vocal communication system of orcas (*Orcinus orca*) has so far been analyzed primarily in terms of the fundamental frequency (F0) modulations, i.e. the frequency of their phonic lips vibration. The calls have been divided into clicks, pulsed calls, whistles and types thereof. By analyzing 61 hours of on-orca acoustic recordings and controlling for the effect of high-frequency components (HFC) and F0, we report structured formant patterns in orca vocalizations including diphthongal trajectories. Broadband spectrogram analysis reveals previously unreported formant patterns that appear independent of F0 and HFC and are hypothesized to result from air sac resonances. This study builds on the recent report of formant structure in vowel- and diphthong-like calls in another cetacean, sperm whales (*Physeter macrocephalus*). Using linguistic techniques, we further demonstrate that some calls are reminiscent of human consonant-vowel sequences, featuring bursts or abrupt decreases in amplitude. We also show that individual sparsely distributed clicks gradually transition into high frequency tonal calls, which aligns with analysis of sperm whale codas as vocalic pulses. The paper makes methodological contributions to the cetacean communication research by analyzing orca vocalizations with both narrowband and broadband spectrograms. The reported patterns are hypothesized to be actively controlled by whales and may carry communicative information. The spectral patterns shown in this study provide an added dimension to the orca communication system that merits further analysis and demonstrates convergent evolutions of similar phonological features in cetaceans (orca and sperm whale) and human communication systems.

## 1 Introduction

Orcas (*Orcinus orca*) are one of the most studied cetacean species with a cosmopolitan distribution across the world oceans and exhibit complex vocalizations and sophisticated social and cultural lives. Orcas are the largest species in the Delphinidae family (with adults ranging in length from 5-9 m) and have the second largest brain in the animal kingdom, after sperm whale. Orcas exhibit several complex social behaviors, such as collaborative birth^1^, animal grief as demonstrated by carrying a dead neonate calf for 17 consecutive days^2^, and self-recognition^3^. As apex predators, orca foraging occurs in relatively shallow epipelagic waters within continental shelves and, as a species, they have been documented to consume over 140 species including fish, cephalopods, marine mammals, seabirds, and marine turtles.^4^ “Residents” are the most studied orca ecotype in the Northeast Pacific, where populations live in stable matrilineal social groups ranging from 5-50 individuals and have specific prey preferences and cultural traits. Residents stays in the same pod, clan and community for their whole life, whether male or female [5, 134]. There are also “transients” who live in the same waters as residents, yet do not interact and have a different preferred prey. This study is based on Northern and Southern resident orcas in the US and Canada^6^.

Unlike some odontocetes that utilize clicks for biosonar and social functions (e.g., sperm whales), orcas produce clicks for echolocation^7–12^ as well as pulsed calls and whistles as social signals^7,13–16^. Orcas’ complex vocalizations^17,18^ also utilize multiple fundamental frequencies simultaneously in biphonated calls, varying their fundamental frequencies, call intensities, and temporal patterns. Orca vocalizations consist of a variety of sounds that have been divided into three main types: clicks, whistles, and pulsed calls^7,13,19^. Clicks are impulsive (*<*1 ms) and highly directional sounds ranging from 10-100 kHz that are used for echolocation, i.e., to navigate and hunt^11^. Whistles are frequency-modulated tonal sounds that can be accompanied by repeated harmonic overtones^7^. These sounds can last several seconds and differ in frequency range, e.g., from 1-74 kHz in the northern Atlantic Ocean^20^. Pulsed calls are rapidly repeated sounds with distinct tonal properties that usually contain abrupt and patterned changes in the pulsing rate^7^ that are variable in length, frequencies, temporal patterns, and volumes. Orcas can also use multiple frequencies simultaneously, as in the case of biphonic calls.^7,21,22^ Orca calls have been clustered into 4-91 call types depending on the study and regions.^7,13,22–26^

This paper adopts the source–filter conceptualization of odontocete vocalizations and the parallel with human vowels recently proposed in Beguš et al.^27^ We treat the fundamental frequency (F0) produced by the phonic lips that generate the *low-frequency component* (LFC) as the source feature. In biphonic calls, a second set of phonic lips produces an additional F0. When this second F0 is higher than the first F0, is called the *high-frequency component* (HFC). An extensive body of work exists on the source features of orca vocalizations^9,18,28^. We analyze formants resulting from resonant frequencies of the air sacs that follow the phonic lips as filter features. To our knowledge, no study thus far have shown a variable formant structure independent of F0 when the effects of the higher fundamental frequency are controlled for. Some attempts have been made to measure formants in orcas^29,30^ to distinguish between individual whales, but the work did not yield meaningful differences and also did not control for biphonation. While orcas have also been shown to imitate human speech, this study focuses on temporal, amplitude, and F0 patterns, and does not measure formants.^31^

In addition to the previously described source features (F0), this study shows that orca vocalizations (in their natural setting) contain filter features in the form of complex and structured formant patterns that have previously been unobserved. We observe these patterns below the HFC, making it unlikely that they are harmonics of the HFC. We further show that these formant features can be independent of F0, and thus likely actively controlled by the whales. The newly identified patterns indicate that orcas orcas can modulate formant structure in consecutive calls, non-consecutive calls, and even within a single call. We exclusively analyze on-whale recordings, where the hydrophone records directly from the whale or from a nearby whale, minimizing the confounds of underwater acoustics and distance-based recording artefacts.

A complex formant structure in toothed whales (odontocetes) was first proposed for sperm whales^27^. Sperm whales vocalize with clicks that they group into *codas* that were previously analyzed almost exclusively in terms of source features (number of clicks and timing). In Beguš et al. (2025,2026)^27,32^, sperm whale codas were shown to be acoustically analogous to human vowels. This study demonstrates that orca vocalizations also possess similar vowel-like features found in sperm whales. In addition, some orca calls are reminiscent of the consonant-vowel sequences found in human languages.

In human language, vowels are sounds of speech with a maximum degree of aperture and high loudness (amplitude). Characteristic of human vowels and consonants is the source-filter production mechanism, where vocal fold vibration (source features or F0) is combined with changes in the vocal tract resonances (filter features or formants).^33,34^ Consonants, on the other hand, are characterized by a higher degree of vocal tract closure. Stops, such as [p], [t], or [k], are produced with full closure of the vocal tract (resulting in a period of silence), followed by a burst^35^ and voice onset time^36,37^. Stops are consonants with the highest degree of closure of the vocal tract. Other consonants, such as nasals, fricatives, or approximants are produced without full closure, but with less opening than vowels. The characteristic of approximants such as [l] or nasals such as [n] is the reduction in the amplitude values and formant structure^38^. The acoustic analysis in this paper suggests that orca vocalizations feature structural analogs to human vowels and some consonants such as stops and approximants (Figure 1).

**Figure 1.**
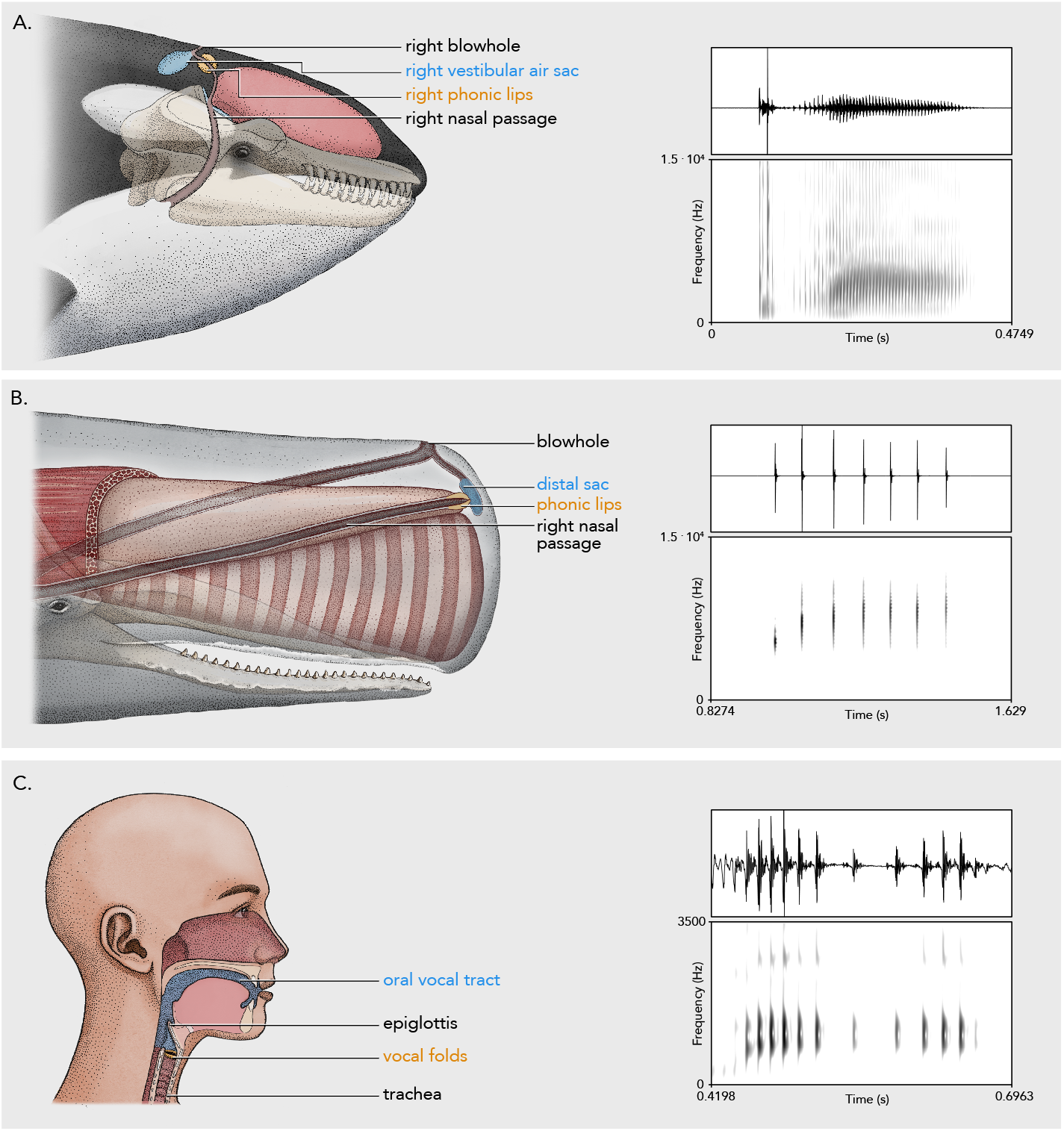
Orca (a), sperm whale (b), and human (c) articulators and vocalizations with waveforms and corresponding spectrograms. All three vocalizations feature formant trajectory. In sperm whales and orcas, the formant begins at a lower frequency and levels off in the second part of the call (a diphthong). In the human syllable, the first part of the syllable likewise shows formant lowering. The human recording is a Mandarin syllable *ma* with the third tone; the third tone results in creaky phonation which illustrates parallels between sperm whale and human vowel system. The orca vocalization is acoustically even closer to human modal phonation during vowel production and additionally features a consonant-like burst and a period reminiscent of voice onset time.

## 2 Data and methods

This study analyzes data from an open source repository of orca vocalizations in Tennessen et al. (2024)^6^. The recordings were made in 2009, 2010, 2011, and 2014^6^ from Northern and Southern Resident orcas in the coastal waters of Washington State, USA and British Columbia, Canada. The recordings were made with D-Tags^39^ placed on whales (as demonstrated in Figure 2). The first author performed a manual acoustic analysis of all recordings from years 2011 and 2014, all recordings of Southern Resident population from 2010, and selected recordings from Northern Residents from 2009 and 2010. A total of 61 hours of D-Tag recordings from the repository were manually analyzed. A total of 1,104 calls or trains of calls were analyzed.

**Figure 2.**
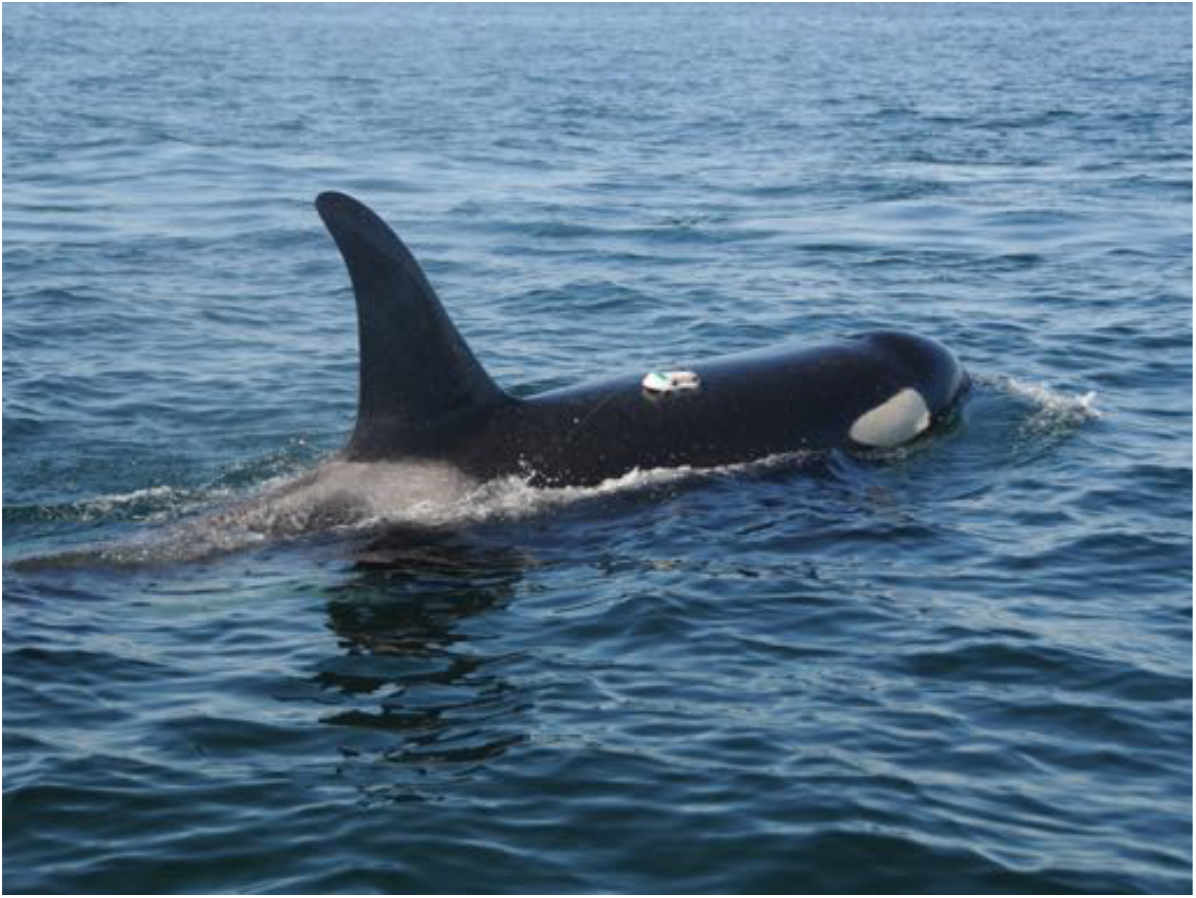
Southern Resident male Orca (K33) with on-whale acoustic sensor in 2010 when the vocalization in Figure S1 was recorded. NOAA NWFSC taken until Permit No. 781–1824.

In this paper, we focus on calls where we can establish an independence between the source features (F0) and formant frequencies and where the formants are not harmonics of HFC. We present waveforms and narrow- and broadband spectrograms that demonstrate different formant profiles under similar F0 contours as well as different F0 contours under similar formant structure. All analyses are accompanied by narrowband spectrograms indicating the presence or absence of the HFC and its value. Because our analyses target the frequency band below the HFC, the reported formant patterns are unlikely to reflect harmonics of the HFC.

We present calls in isolation that are plausibly produced by the focal whale based on amplitude values. While amplitude values alone cannot guarantee that the analyzed vocalizations are produced by the focal whales, high-amplitude signals are suggestive of either the focal whale or a nearby whale which minimizes underwater acoustic disturbances or artifacts. Not all calls presented here are focal, but they are all of high amplitude such that the possibility of acoustic artifacts is substantially reduced.

All spectral analyses were performed in Praat^40^ with standard settings for spectrogram creation. The spectrograms were created with window size ranging from 0.7 ms and 2 ms (specified for each spectrogram) for broadband spectrograms and 10 ms for narrowband spectrograms. Dynamic ranges were set at different levels from approximately 20 dB to 40 dB. The frequency range is 0–15,000 Hz unless specified differently. Spectrograms were produced with Gaussian windowing, 1,000 time steps and 250 frequency steps, autoscaling, and 6.0 dB/octave pre-emphasis. Spectra were created with Praat’s *Spectrum* function which performs the Fast Fourier Transform on the selected time series data.^40^ Pitch tracks were obtained with the raw autocorrelation method with various pitch floor and ceiling values (as indicated in the graphs): silence threshold was set to 0.03, voicing threshold to 0.45, octave cost to 0.01, octave jump cost to 0.35 and voiced/unvoiced cost to 0.14 (Praat standards).

## 3 Results

We demonstrate a clear formant structure in orca tag recordings. We show that this formant structure can be independent of F0 (Section 3.1). We also describe diphthongal (Section 3.1.3) and consonantal patterns (Section 3.1.4) in orca vocalization and argue that the difference between individual non-echolocation clicks, pulsed calls, and high frequency tonal calls largely reflects differences in pulse rate (F0), i.e. the spacing between successive pulses (Section 3.2). This links orca vocalization especially closely to the coda vowels of sperm whales (Section 3.2). Based on the spectral measurements, Section 3.3 proposes a potential articulatory mechanism for the described spectral patterns.

### 3.1 Formant structure

Here, we present broadband and narrowband spectrograms of different calls with the same F0 structure but different formant structure (Section 3.1.1 and Figures 3 and 4). We also show that a trajectory in formant structure is possible within a single call. This can happen in calls with substantially different F0 trajectories (Figure 5(left)). Additionally, we show that a single call with level F0 can have variable formant structure (Figures 5 left and 6). Finally, we show that a rising-falling formant trajectory is possible in a call with only a rising F0 pattern, which suggests that the F0 trajectory does not crucially affect formant trajectories (Section 3.1.3; Figure 7). Together, these results suggest that the formant patterns (filter features) are independent of F0 patterns (source features). The observed formant patterns fall within orcas’ hearing range^41,42^. We argue that calls previously analyzed as a single call type are spectrally different when formant frequencies are taken into account.

**Figure 3.**
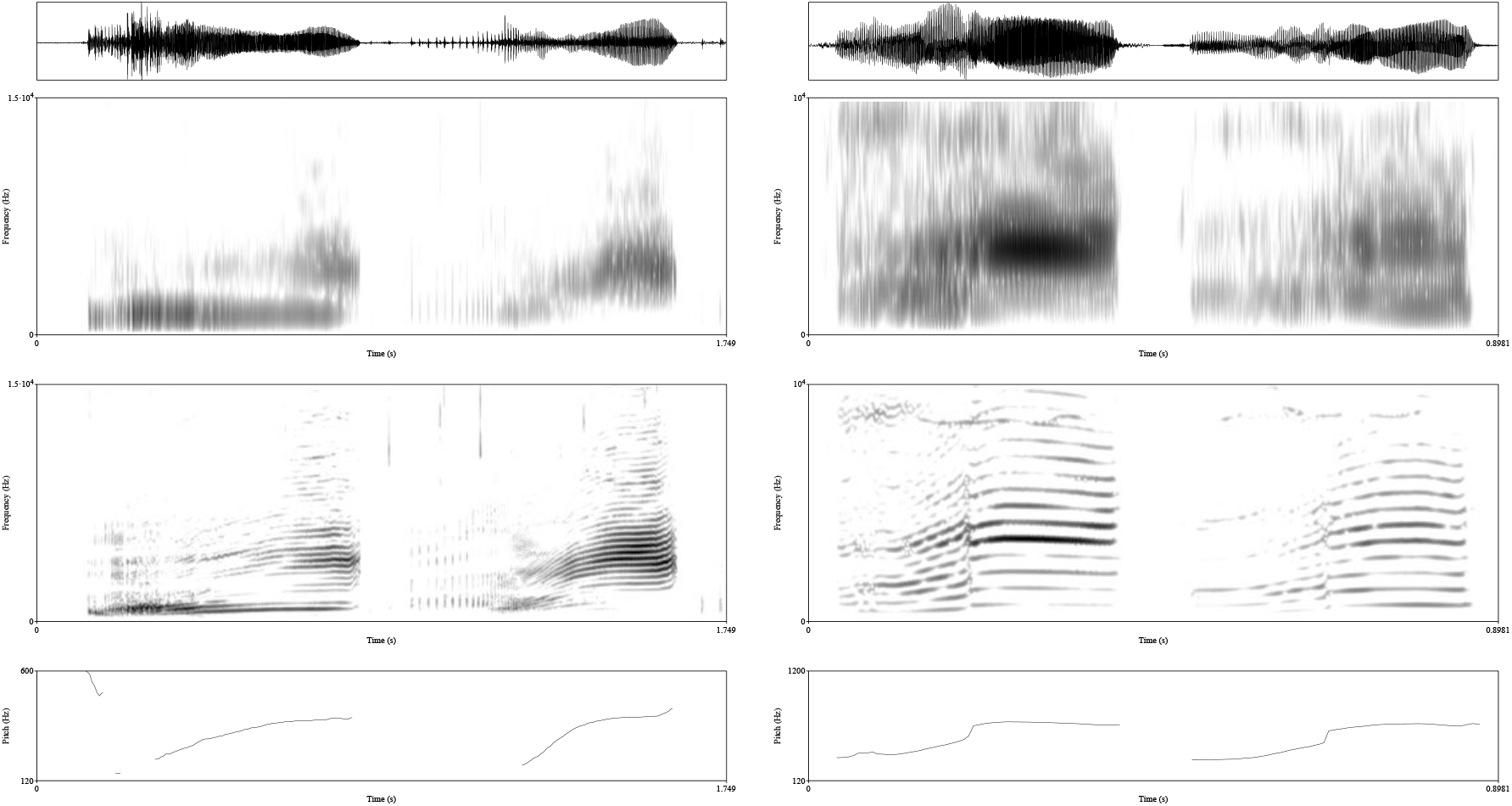
(**left**) Waveforms, a broadband (1 ms window), narrowband spectrogram (10 ms window; both 0–15,000 Hz), and F0 contour (raw autocorrelation) of two consecutive calls (with 0.65 s of silence removed between them) recorded when a 19-year old female (L91) from the Southern Resident population was wearing a tag. (**right**) Waveforms, a broadband (1 ms window), narrowband spectrogram (10 ms window; both 0–10,000 Hz), and F0 contour (raw autocorrelation) of two calls recorded when the same 19-year old female (L91) from the Southern Resident population was wearing a tag. The calls were recorded on the same tag approximately 160 s apart whereby the first call was uttered after the second call.

**Figure 4.**
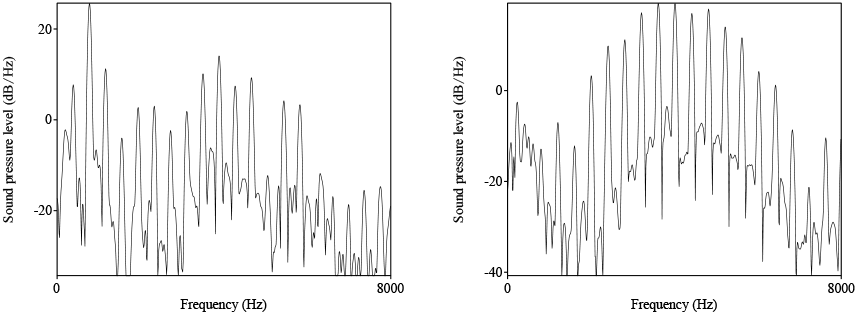
Spectra (0–8,000 Hz) of the 0.05 s window of the two calls in Figure 3 (left). The two figures show clear differences in spectral profiles of the two calls.

**Figure 5.**
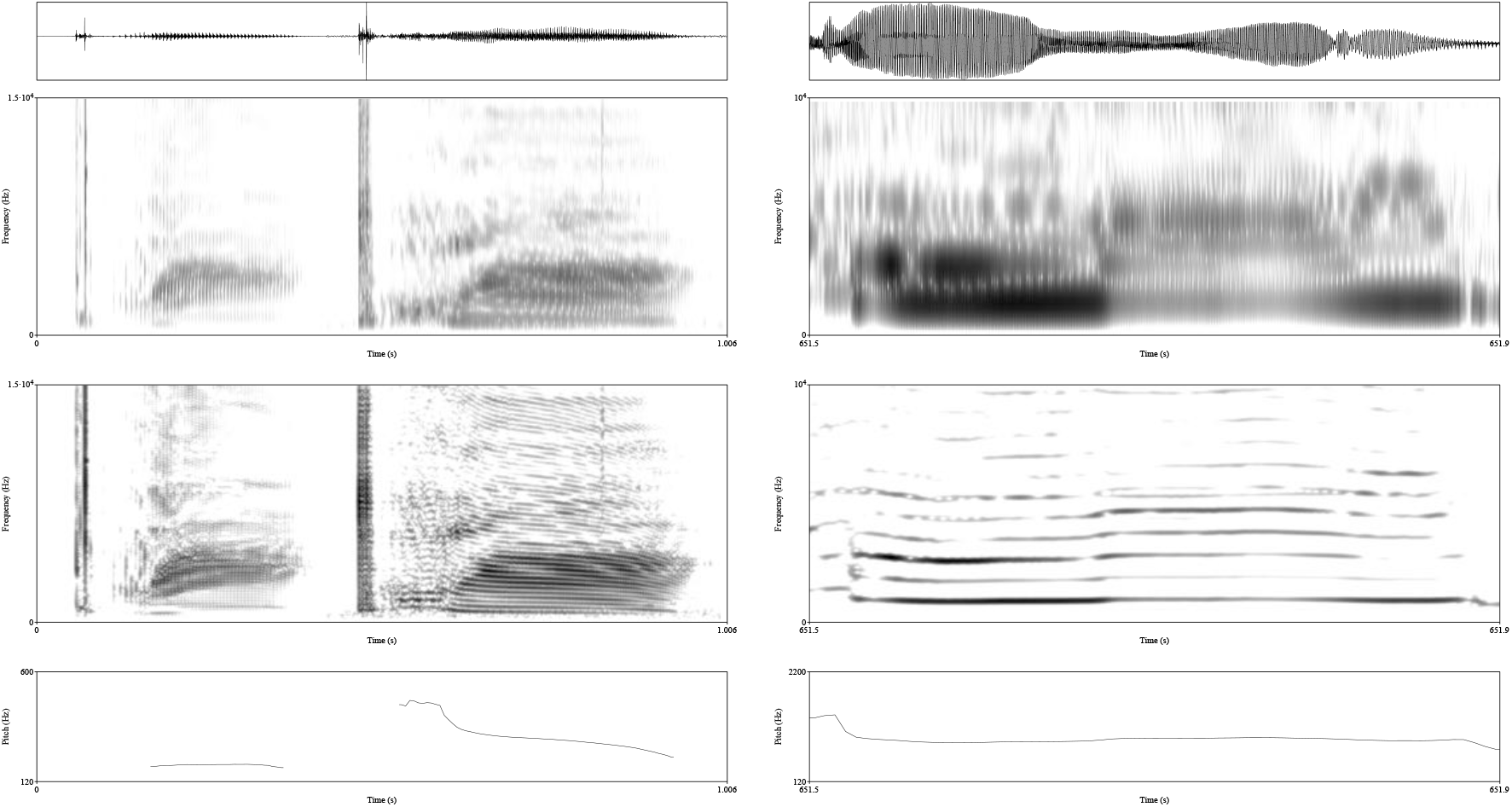
(**left**) Waveforms, a broadband (2 ms window), narrowband spectrogram (10 ms window; both 0–15,000 Hz), and F0 contour of two calls with burst-like onset and diphthongal formant patterns, recorded when a 6-year old male from the Northern Resident population was wearing a tag. (**right**) Waveforms, a broadband (1 ms window), narrowband spectrogram (10 ms window; both 0–10,000 Hz), and F0 contour of a single cropped call recorded when a 19-year old female (L91) from the Southern Resident population was wearing a tag.

#### 3.1.1 Consecutive calls

Figure 3(left) shows two consecutive calls with a similar F0 trajectory. Despite this similarity, the calls exhibit markedly different formant structure, especially in the second half of each call. The narrowband spectrogram suggests that both calls are produced with a single set of phonic lips, as only one harmonic series is visible and no HFC is present. Spectra with a 0.05 s window in Figure 4, taken from the final part of each call where the differences are most pronounced, show substantially distinct profiles of resonant frequencies. The first call shows two formants (with the first formant not coinciding with the first harmonic, as is evident from Figure 4), whereas the second call contains only a single formant.

Another case of a variable formant structure despite the same F0 profile is given in a call in Figure 3 (right). Each call can be divided into two parts, a first half with a relatively lower F0 and a second half with a relatively higher F0. Based on the fundamental frequency, the two calls are nearly identical and would be treated as repetition. However, the first call has a clearly pronounced single formant, whereas the second call has two. The HFC is above 8 kHz in both calls, which means that the formant structure below this threshold is not likely caused by HFC.

Sperm whales also produce codas with one or two formants. The presence of one versus two formants within the relevant frequency range in sperm whale codas (as is the case here for the difference between one or two formants within a frequency range) is the primary distinction between the *a*-coda vowel and the *i*-coda vowel^27^. The orca calls analyzed here therefore provide a close parallel to the spectral properties observed in sperm whales.

#### 3.1.2 Single calls

Complex formant structure independent of F0 of HFC can also occur within a single call. Figure 5 (right) shows spectrograms of a single call recorded from a tag on a Southern Resident orca. The entire call has a flat F0 trajectory from the low-frequency component, but clear differences in formant structure. The call can be divided into three parts based on formant structure. The HFC begins at about 5 kHz—structure below this threshold thus stems solely from one set of phonic lips and their resonant frequencies. The first quarter of the spectrogram shows two formants that gradually transition into a single formant below 5 kHz. The last part of the call also has a single formant below 4 kHz with a peculiar arc-shaped second formant. The middle part of the call has a more distributed spectral profile. What previously appeared as a simple call with a flat spectral profile received a complex and variable formant structure under our proposed analysis. A reverse pattern to this call is observed in a different deployment. Figure S1 (Supplementary Figures) shows a call with a structure similar to the call in Figure 5 (right) but in the opposite sequence: the single-formant part is ordered first, followed by a period of decreased amplitude and distributed spectral profile, followed by a period where a single formant transitions into a period with clearly pronounced two-formant structure. The calls described so far share similar F0 trajectories but differ in formant structure. To demonstrate that F0 and formant structure vary independently, we also analyze the converse pattern: calls that share similar formant structure but differ in F0 trajectories. Figure 5 features two calls with substantially different F0 trajectories, but with similar diphthongal pattern in which the first part of the call starts with a lower formant and the second part levels off at a higher frequency.

#### 3.1.3 Diphthongs

Diphthongs are defined as cetacean calls with substantial formant trajectories^27^. Human speech features diphthongs such as English [aI] (as in *rise*). Structured and repetitive diphthongal patterns realized as formant trajectories have also been observed in sperm whales^27^. These trajectories occur across individuals and have been shown to be independent of whale movement. Here, we report comparable formant trajectories in orca calls. Figure 5(left) shows two calls with similar and pronounced formant trajectories. In both, the first formant (F1) starts low, rises rapidly, and then levels off for the remainder of the call. The two calls have substantially different F0 trajectories and absolute F0 values, which is consistent with the independence of F0 and formant production.

To estimate the magnitude of formant movement during the diphthongal calls, spectra of individual pulses of the first call in Figure 5 were measured with Praat’s^40^ *Spectrum* function. The spectral peak of the first pulse with a clear formant structure is 2,527 Hz, whereas a pulse in the middle of the call has a spectral peak of 3,736 Hz. Figure 6 shows the spectra of the two pulses illustrating an upward trajectory.

**Figure 6.**
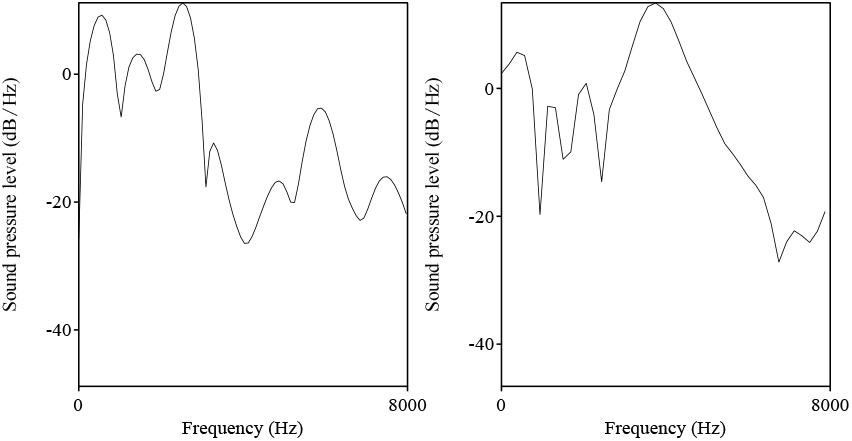
Spectra of two pulses, one at the beginning and one in the middle of the first call in Figure 5 (left), recorded when a 6-year old male from the Northern Resident population was wearing a tag.

**Figure 7.**
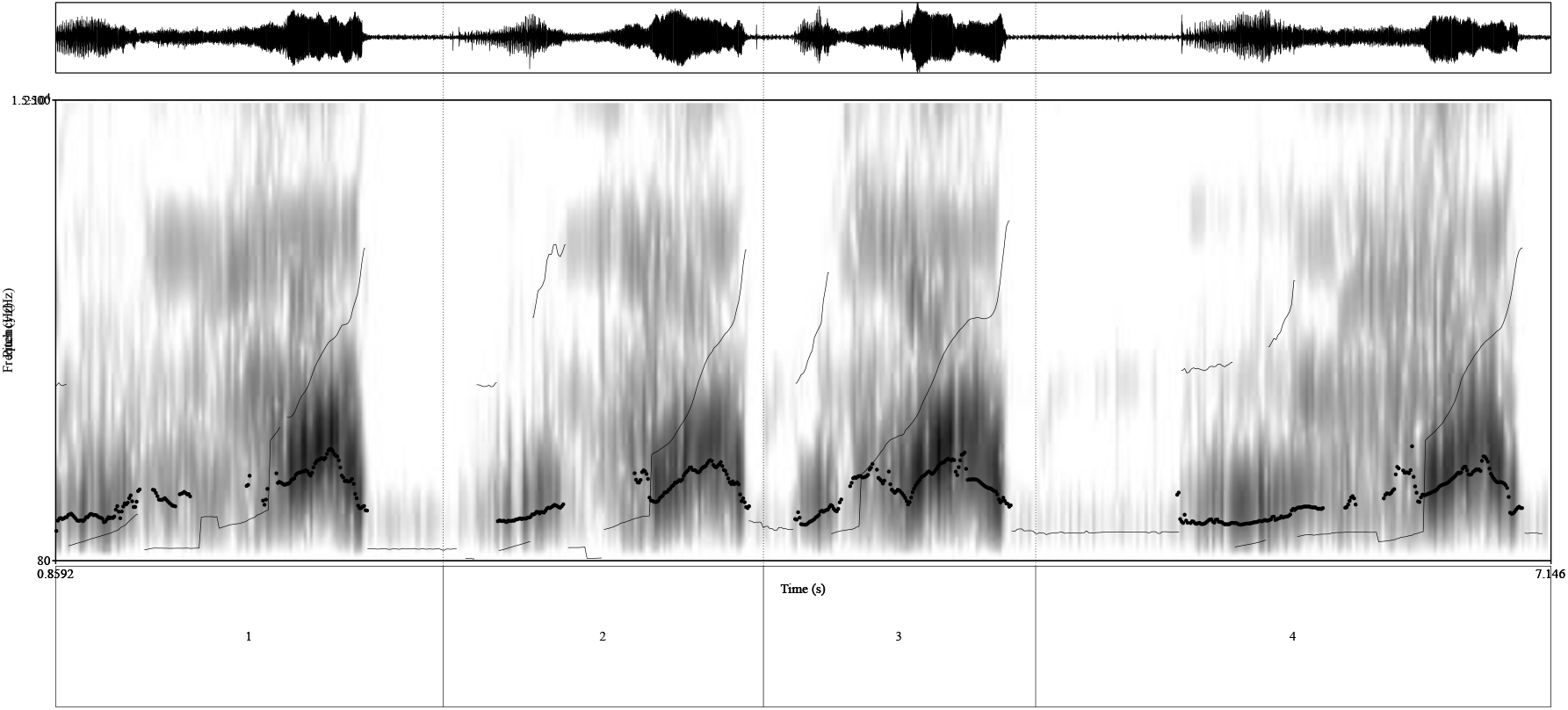
Waveforms, a broadband (0.7 ms window, 0–15,000 Hz), F1 contour (speckled) and F0 contour (lines; raw autocorrelation) of four calls were uttered within a time span of 30 seconds, recorded when a 24-year old female (L72) from the Southern Resident population was wearing an acoustic sensor. The narrowband spectrogram is given in Supplemental Figure S3.

That the described diphthongal trajectories are not an artifact of F0 trajectories is strongly suggested by four calls in close proximity in Figure 7. We show spectrograms, formant tracks, and pitch tracks of the four calls, which come from the same tag. The last part of each call features a clear rise in F0 in all four calls, and an intriguing formant transition. The F1 pattern in the last part of the call first rises and then falls, for a clear rising-falling pattern. This arch-like formant movement happens despite the consistent rise in F0. Given that the call is repeated four times with the exact same spectral structure suggests that it is likely controlled by the whales. The HFC is well above the observed rising-falling F0 contour (as clear from Figure S3).

#### 3.1.4 Consonant-like structure

In addition to formant structure described so far that parallels sperm whale coda vowels and human vowel patterns, we observe several calls of the type illustrated in Figure 5(left). All of these calls start with a few pulses that appear distinct from the rest of the call. The main tonal pulses of the call start with some latency after the initial few pulses. The pulses are of high-intensity and their frequencies are relatively evenly distributed as the spectrograms in Figure 5(left) suggest. Acoustically, these calls are the closest approximation of the consonant+vowel sequences found in human language, with the first few clicks acting acoustically as bursts in human language. The orca “bursts” are not present in every call–a call can start with or without these bursts (as is clear from other waveforms and spectrograms in Figures 3, 5). In human speech, some syllables have consonant onsets and others don’t. As already mentioned, there is a latency between the bursts and the full onset of the periodic high frequency tonal pulsed call that follows, which is highly reminiscent of the voice onset time in human speech.

Another call type with a clear burst-like onset followed by a level formant with no formant trajectories is illustrated in Figure S4 (Supplementary Figures). Some calls also contain periods of abrupt amplitude reduction, reminiscent of human sonorant consonants. Figure 8 shows a call that begins with a vowel-like segment exhibiting a single prominent formant, followed by an abrupt decrease in amplitude and loss of clear formant structure. Approximately halfway through the call, a vowel-like formant structure, this time with two formants, starts appearing. The call ends with irregular vibration and increased aperiodic energy reminiscent of frication-like noise in human speech. The spectral analysis thus suggests that orca calls can have a flat F0 structure but clearly delineated periods of differentiated formant structure.

**Figure 8.**
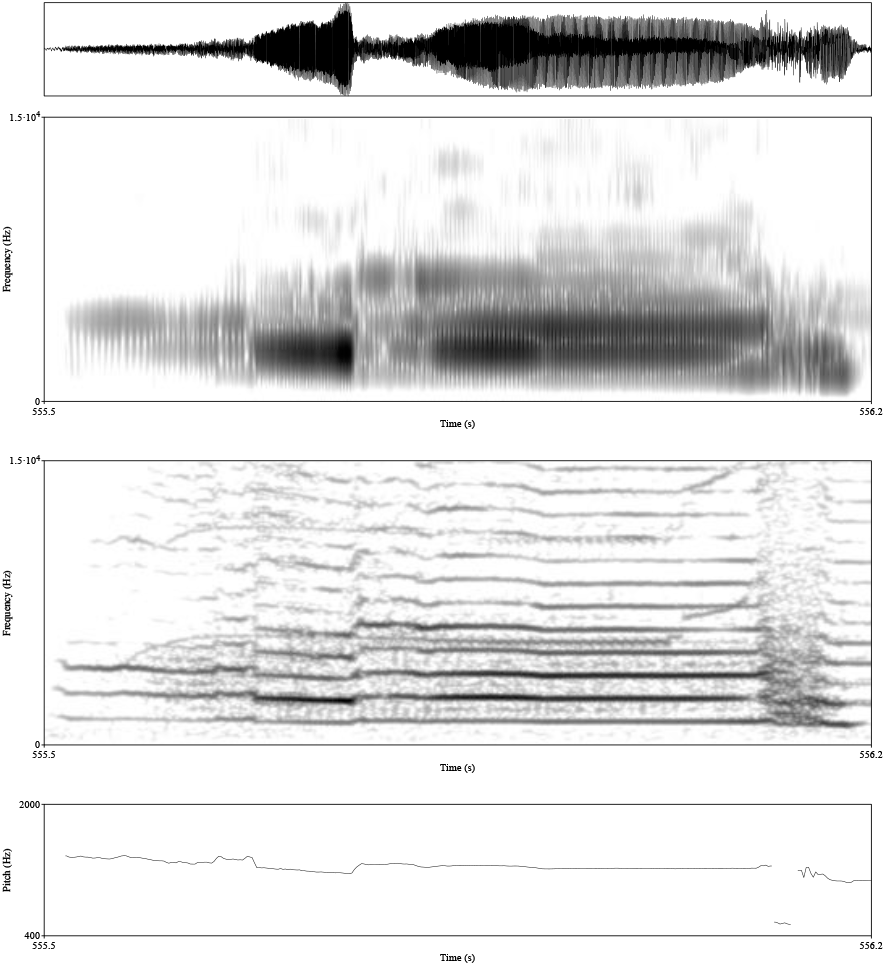
Waveforms, a broadband (1 ms window), narrowband spectrogram (10 ms window; both 0–15,000 Hz), and F0 contour of a call with consonant-like abrupt change in amplitude recorded when a 9-year old male (K33) from the Southern Resident population was wearing an acoustic sensor.

These consonant-like acoustic properties could, in principle, be used by orcas to encode contrast in meaning. At present, however, these resemblances are only acoustic: we do not yet have a firm understanding of their function or articulatory mechanisms. Even so, the observed patterns suggest that orca vocalizations might be highly complex, and that this complexity may be acoustically similar to human speech.

### 3.2 Clicks and pulsed calls

In Beguš et al. (2025, 2026)^27,32^, we show that sperm whale clicks act as vocalic pulses, and we hypothesize that sperm whale codas (groups of clicks) function analogously to human vowels: traditional coda type parallels source features in human vowels (F0), while formant structure results from resonant frequencies. Under this view, sperm whale codas have a much lower F0 than human vowels, but are otherwise acoustically analogous.

Orca vocalizations provide additional support for this framework. In Figure 9, two calls gradually transition from what would be analyzed as individual, non-echolocation clicks into a high-frequency tonal call. The initial part of pulsed call is acoustically closest to sperm whales’ codas. Codas appear as pulsed calls because they are slower. Sperm whales transition from individual clicks into tonal calls only rarely (e.g. in trumpet calls^43^). One possibility is that differences in sound-production anatomy (e.g., phonic-lip morphology) constrain the range of click-to-tone transitions across species. Orcas with smaller phonic lips can produce both coda-like calls as well as higher frequency pulsed calls.

**Figure 9.**
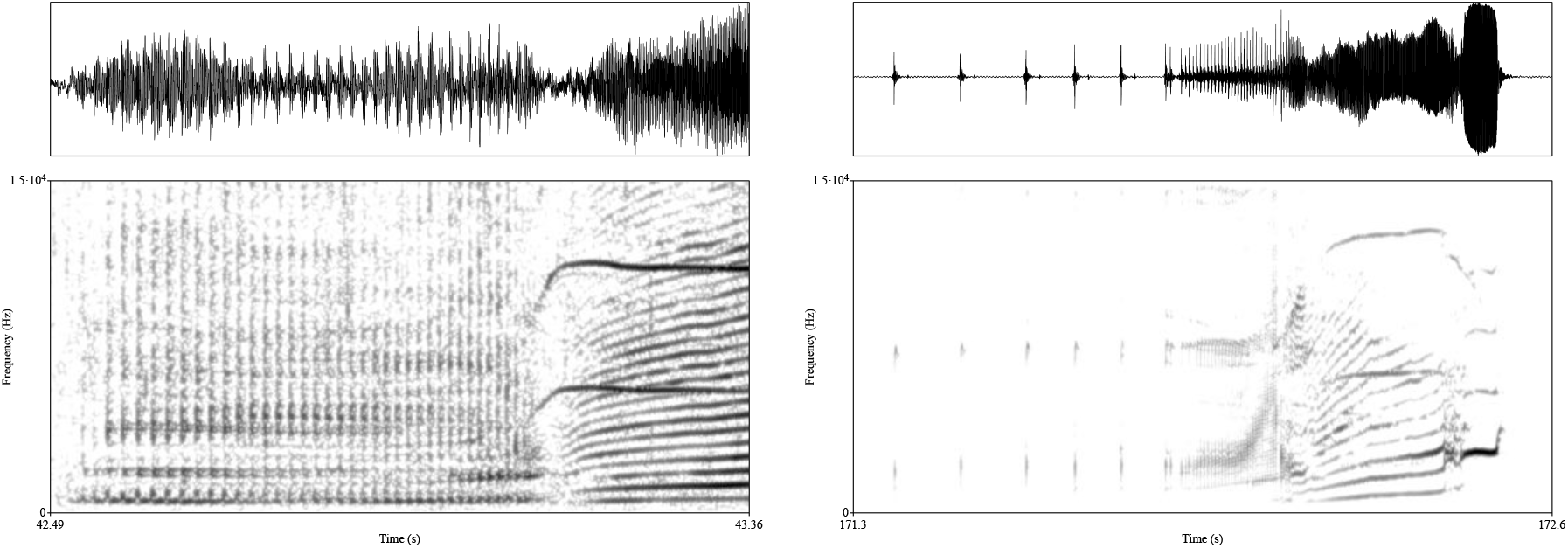
Waveforms, a narrowband spectrogram (10 ms window; both 0–15,000 Hz) of two click train calls that gradually transition into a high frequency call as F0 increases. The call on the left was recorded when a 23-year old male (L85) from the Southern Resident population was wearing an acoustic sensor. The call on the right was recorded when a 24-year old female (L72) from the Southern Resident population was wearing an acoustic sensor.

It is possible that some acoustic differences between low frequency clicks and tonal pulses remain as the tissue or muscular tension in phonic lips during slow pulsed calls could differ from that during higher-frequency tonal call production. This is comparable to human “creaky” voice, where creaky phonation affects the acoustics of individual pulses^44–47^. Crucially, the gradual transition from a train of clicks/pulsed call to a high frequency vowel-like tonal call that primarily depends on F0 rate is precisely what we propose for the analogy between human vowels and sperm whale vowels: codas are understood as very low frequency vowels, acoustically similar to creaky phonation in humans^32^. Figure 9 further shows that orca low frequency pulsed calls exhibit discernible formant structure as well.

In Beguš et al. (2026)^32^, we hypothesize that whales have a lower F0 threshold at which pulse trains are perceived as pitch rather than as discrete beats. In humans, this threshold is around 30 Hz^48–50^. If sperm whale clicks are analyzed as vocalic pulses, the base F0 of sperm whale codas is approximately 4-20 Hz^32^. Along these lines, orcas might exhibit a broader span of pitch perception, potentially extending from low-frequency click pulses to high-frequency tonal calls. This framing is consistent with the range of orca hearing^41,42^.

### 3.3 Possible articulatory mechanism

In Section 3.1, we observe multiple formant patterns in the frequency range from 0 Hz up to the F0 of HFC. We hypothesize that these formant patterns result from changes in the articulatory configuration inside orcas’ nasal complex. Orcas have a complex set of air sacs and passages that could plausibly cause the observed formant patterns. One possible resonator is the vestibular air sac with its associated passages, given that airflow enters the vestibular air sac directly after passing the phonic lips. We use a simplified tube model to estimate the effect of these potential resonators.

In earlier work on sperm whales^27^, we proposed that resonator size can be approximated using the standard tube-model relationship between resonant frequencies and tube length (Eq. 1^34^).

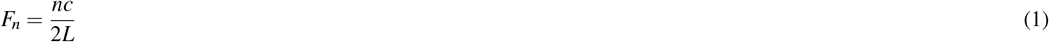

To estimate the resonator length change required to produce the first formant trajectory in the first call in Figure 3(right), we measure the formant peak of a single pulse near the beginning of the call (with a window of 15 ms) and again near the middle of the call. The formant peak of the pulse at the beginning of the call is measured at 2,527 Hz whereas the formant peak at the middle is measured at 3,736 Hz (see also Section 3.1.3 above). Assuming that the resonant body in orcas can be approximated as a tube closed at both ends, the length of the tube is calculated to be 6.8 cm (according to Eq. 1) given that F1 is measured at 2,527 Hz near the beginning of the call. The length of the tube is calculated to be 4.6 cm near the middle of the call given that the first resonant frequency is measured at 3,736 Hz (assuming 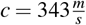for both).

If the resonator is approximated as a tube open at one end, the corresponding lengths of the tube (L) predicted by Eq. 2 are cm near the beginning of the call and 2.3 cm near the middle.

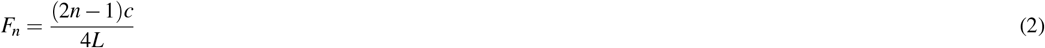

These calculated differences in tube length are well within the plausible range of the vestibular air sac size. The vestibular air sac receives airflow after it passes the phonic lips^18^. Estimating the size of these air sacs is difficult. Mead (1975)^51^ reports the right vestibular sac to be larger in size than the left with the right “anterior fold” measuring 2 cm. It is therefore reasonable to assume that orcas could manipulate the vestibular air sacs such that the resulting acoustic tube length changes by 2.2 cm or 1.1 cm, respectively (depending on whether the tube is modeled as closed at both ends or open at one end).

## 4 Discussion and conclusion

Our acoustic analysis suggests that orca vocalizations exhibit complex formant structure that is likely independent of F0 and actively controlled by the whales. We describe several previously unobserved formant patterns and hypothesize that these formants result from the resonant bodies inside the nasal complex. We minimize the confounds of underwater acoustic artifacts by exclusively analyzing on-whale acoustic sensor recordings. By combining broadband and narrow-band spectral analyses, we control for the effect of biphonation in calls showing that the formant structure does not result from the fundamental frequency (F0) of the HFC. By showing that calls with constant F0 can have different spectral patterns and conversely, calls with similar spectral patterns can have diverse F0 contours, we conclude that F0 does not crucially affect the observed formant patterns.

This study builds upon reports of vowel-like patterns in sperm whales Beguš et al. (2025, 2026)^27,32^ and suggests timing is the crucial feature in the study and analysis of cetacean vocalizations. For example, sperm whales’ clicks in codas were sped up for vowel-like structures to become evident to human researchers.^27^ Alternatively, orca calls in this study were slowed down for their vocalic features to be discerned. For the discovery of coda vowels, the crucial step was to remove space between the clicks; for the discovery of formants in orcas, the window size was decreased for spectral analysis, as their calls are of higher frequency. Here we decreased the analysis window of orcas’ calls in line with this shift of timing, to appropriately analyze calls in terms of narrow- and broadband spectrograms.

In addition to vocalic features, human speech includes characterized by a decreased aperture of the vocal tract. So far, consonant-like sound have been documented primarily in primates^53,54^. Here, we show that orcas also produce consonant-like elements. The newly described consonants in orcas are reminiscent of stop consonants and nasal/approximant consonants in humans. While the exact mechanism of burst production in orca remains unknown, we speculate that these bursts are produced with phonic lips. In this respects, the observed consonant-likes sounds are acoustically parallel to ejective consonants in human speech where the burst is produced by a release of the vocal cords (i.e. the same source that produces vocalic pulses). Our spectral analysis also suggests that calls with seemingly flat F0 structure contain periods of differentiated formant structure with clearly pronounced segment boundaries that apear combinatorial and reminiscent of human consonant and vowel sequences (Figure 5 right, 8).

Taken together, orca vocalizations (along with sperm whales) represent one of the highly complex systems in the animal kingdom. Figure 10 summarizes parallels between sperm whale, orca, and human speech. Across all three systems, the source of phonation is vocalic pulses, which can be regularly periodic (modal phonation in humans, high frequency tonal calls in orcas) or more widely spaced (creaky phonation in humans, pulsed calls in orcas, or codas in sperm whales). In orcas, pulsed calls can gradually transition into tonal calls. All three systems thus include vocalic pulses as source features. In humans, they are produced with vocal cords, in orcas and sperm whales with phonic lips (Figure 1). All three systems also exhibit filter features: formant patterns with one or more formants distributed across the spectrum, typically below 15000 Hz. We call these vowel features and speculate that in sperm whales and orcas they arise from resonances of air sacs. All three systems also feature diphthongs, where pulses feature formant trajectories. Humans and orcas are hypothesized to share consonantal features as well. Figure 10 also illustrates that the high-frequency component is only present in orcas, as neither humans nor sperm whales have two sets of phonic lips.

**Figure 10.**
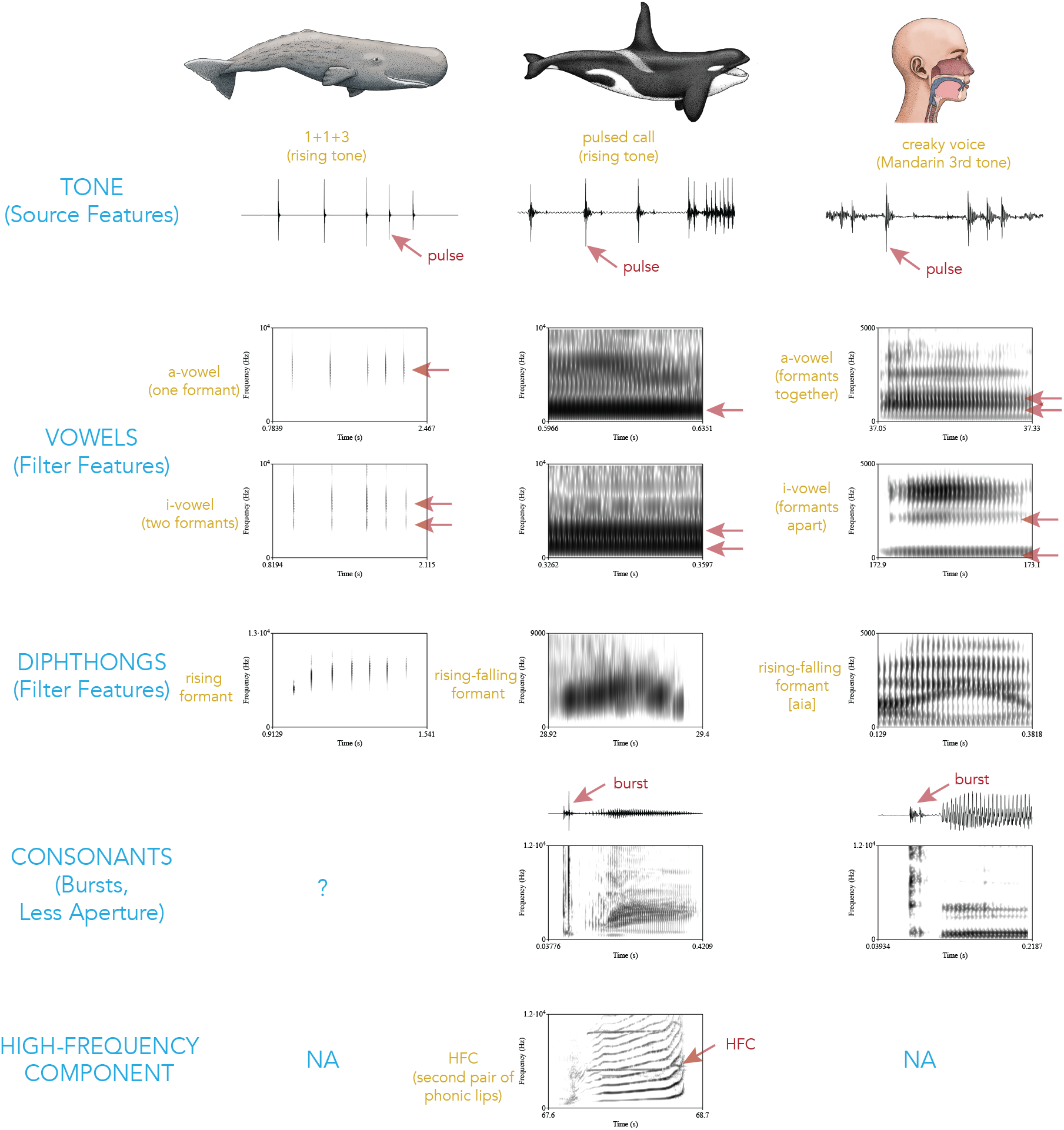
A summary of parallels between sperm whale, orca, and human speech. The human vowel illustration is recorded by a male Mandarin and Georgian speakers. The human consonant is an ejective consonant uttered by a female Georgian speaker^52^. Sperm whale vowels are from Beguš et al.^27^. Orca vocalizations are from figures in this study.

In *Historia Animalium* (Book IV.9), Aristotle writes that dolphins have voice or phonation, sometimes translated as vowel-like sounds (ἔστι γὰρ τούτῳ φωνή). He continues, however, that they don’t have a loose tongue (γλῶτταν οὐκ ἀπολελυμένην) or lips^55^ (χείλη) in order to produce articulate (consonantal) sounds^55^ (ἄρθρον τι τῆς φωνῆς ποιεῖν). However, it is now possible that *Orcinus orca*, a member of Delphinidae, is capable of complex consonantal sounds despite not producing them with their lips or tongue, but rather with articulators within the nasal complex.

These results also have potential implications for the protection of orcas from anthropogenic noise. Orcas are known to adjust their communication in response to presence of noise (the so-called Lombard effect)^56^. Formant structure described in this paper may also be prone to noise interferences and could represent an additional dimension for understanding the impacts of anthropogenic noise^57^.

As orcas are the second cetacean species to exhibit vowel-like features, this provides further evidence that odontocetes may control their articulators in ways that generate formant structure, and that such vocalic (and consonantal) patterns may have communicative function. More broadly, this study suggests that researching the vocalizations of other species of cetaceans with the proposed approach, especially among the 78 species^58^ of extant odontocetes, may reveal similar patterns.

Given that structured formants have been found in both sperm whales^27,32^ and now orcas, this study also provides methodological tools for finding formants in the vocalizations of other toothed whales. We hypothesize that other odontocetes have sufficient flexibility and complexity of air sacs within their nasal complexes to produce controlled resonances analogous to those found in sperm whales and now orcas. Methodologically, our study underscores that each call should be analyzed using broadband and narrowband spectrograms. Appropriate window lengths for narrow- and broadband spectrograms will vary across species and call types, and should be selected with reference to the F0 of the low-frequency component. If variable formant structure is observed within individuals across different F0 contours in additional species, this would support the hypothesis that formant frequencies constitute a potentially meaningful feature of odontocete communication.

The presence of structured formant patterns in at least two species of odontocetes provides a basis for the study of evolution of complex learned vocalizations. Whales and humans are both placental mammals who share physical and physiological characteristics, having a last common ancestor in Boreoeutheria about 92 million years ago^59^. Understanding the parallel development of these complex systems might shed light on the evolutionary pathways of human and non-human learned vocalizations.

## Data Availability

These data are collected by NOAA Fisheries’ Northwest Fisheries Science Center, and Fisheries and Oceans Canada and are available: https://zenodo.org/records/13333019. For animal ethics statements, see Tennessen et al. (2024).^6^

## Author Contributions

G.B. noticed the spectral patterns and analyzed the data. G.B. and D.F.G. wrote the manuscript. D.F.G. obtained project funding. M.H. and B.W. provided the data and metadata and participated in the drafting of the manuscript.

## Acknowledgments

This study was funded by Project CETI via grants from Dalio Philanthropies and Ocean X; Sea Grape Foundation; Rosamund Zander/Hansjorg Wyss through The Audacious Project: a collaborative funding initiative housed at TED. We thank Jennifer Tennessen for open-sourcing the on-whale bioacoustic data which served as the basis for this analysis, and members of the Project CETI team (Ronald Sprouse, Shane Gero, and Giovanni Petri) for their feedback.

## Competing Interests

The authors have no competing interests to declare.

## A Supplemental Figures

**Figure S1.**
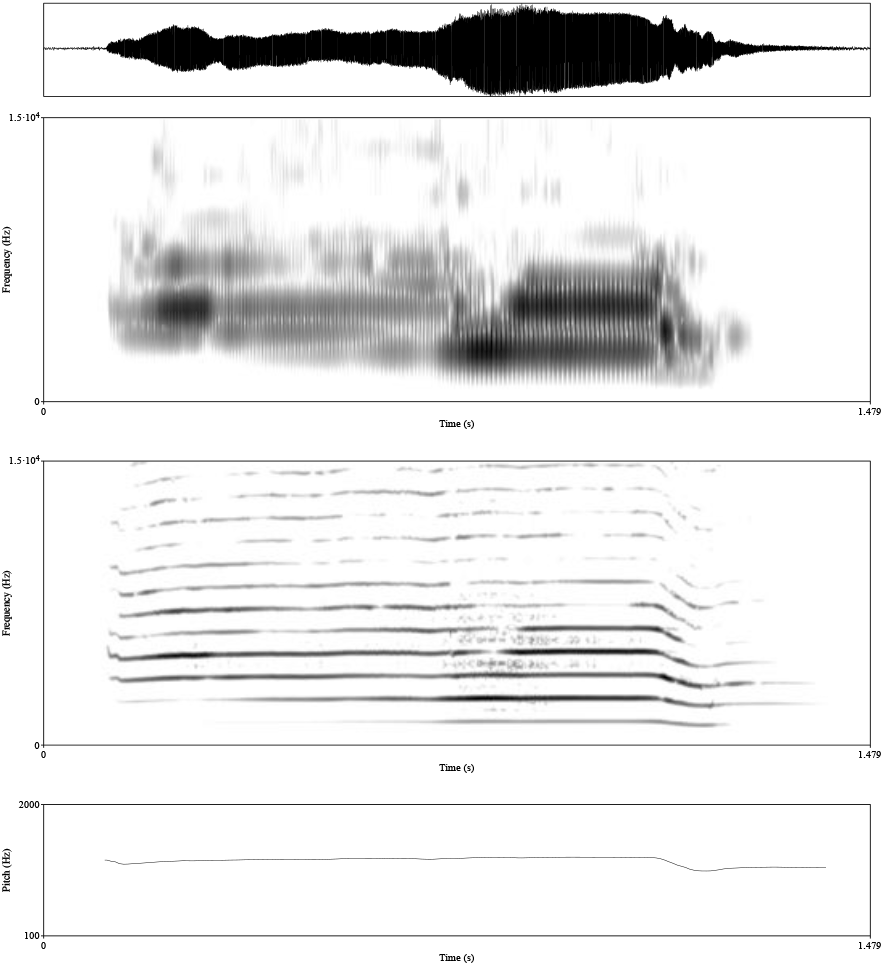
Waveforms, a broadband (1 ms window), narrowband spectrogram (10 ms window; both 0–15,000 Hz), and F0 contour (raw autocorrelation) of a call with an ordering of the structure opposite to the call in Figure 5 (right), recorded when a 9-year old male (K33) from the Southern Resident population was wearing an acoustic sensor.

**Figure S2.**
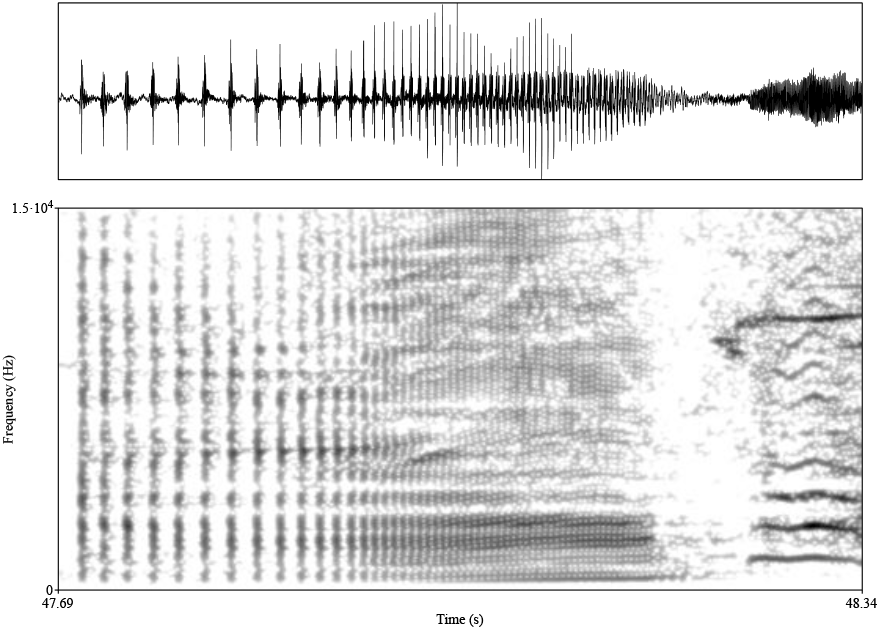
Waveforms, a narrowband spectrogram (10 ms window; 0–15,000 Hz) of a click train that gradually transitions into a vocalic-like high frequency tonal call as F0 increases, recorded when a 23-year old male (L85) from the Southern Resident population was wearing an acoustic sensor. The initial part of the call is highly reminiscent of sperm whale codas.

**Figure S3.**
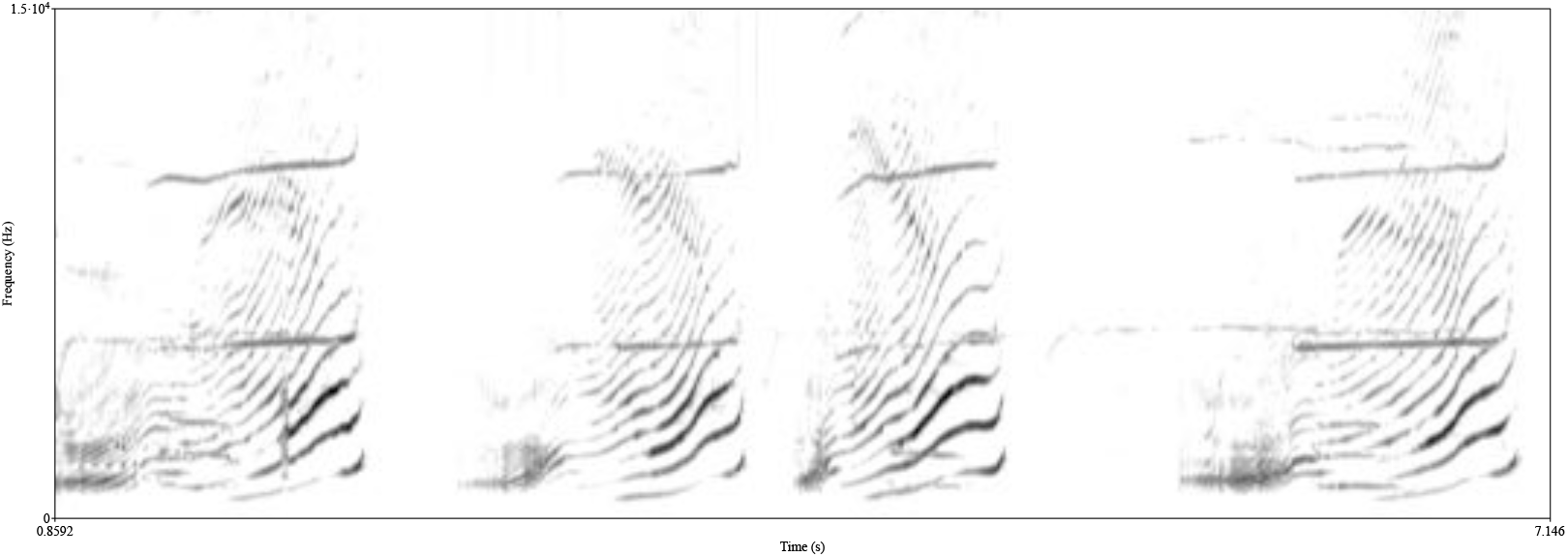
Narrowband spectrogram (10ms window) for the sounds in Figure 7.

**Figure S4.**
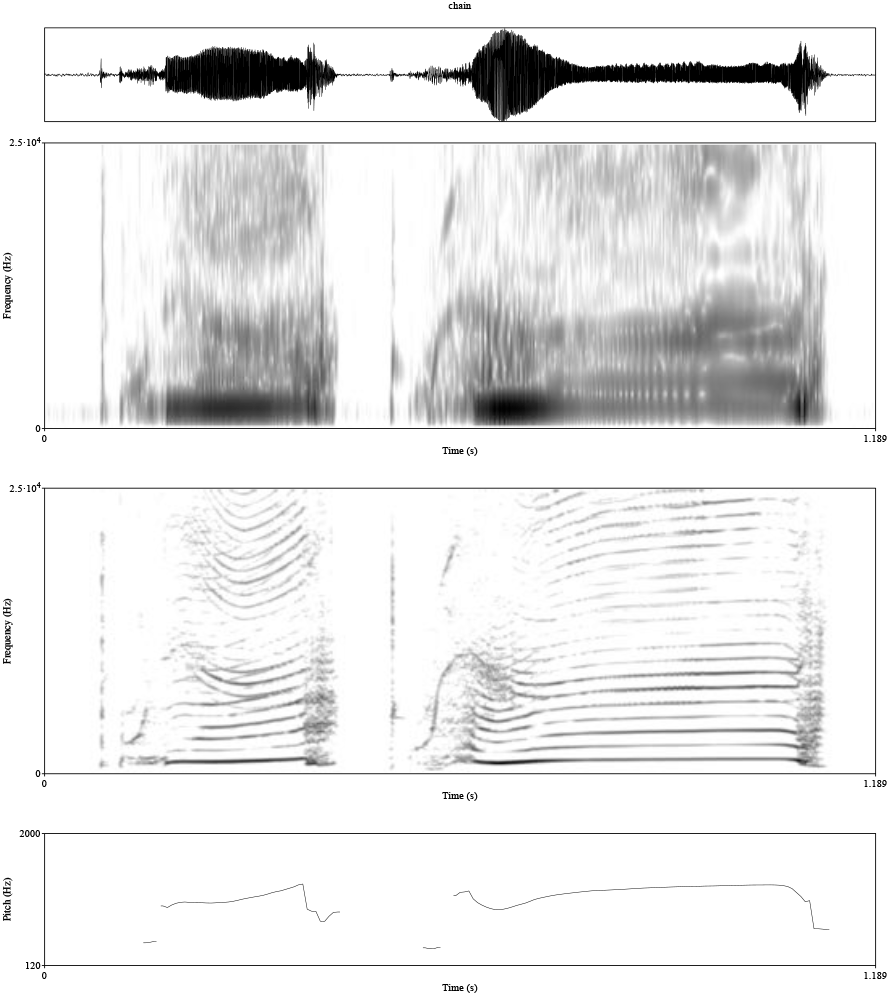
Waveforms, a broadband (0.7 ms window), narrowband spectrogram (10 ms window; both 0–25,000 Hz), and F0 contour (raw autocorrelation) of two calls with a clear burst-like element at the beginning of the call, recorded when a 12-year old female from the Northern Resident population was wearing an acoustic sensor.

## Notes

### Competing Interest Statement

The authors have declared no competing interest.

